# In the murine and bovine maternal mammary gland signal transducer and activator of transcription 3 is activated in clusters of epithelial cells around the day of birth

**DOI:** 10.1101/2023.11.22.568246

**Authors:** Laura J. A. Hardwick, Benjamin P. Davies, Sara Pensa, Maedee Burge-Rogers, Claire Davies, André Figueiredo Baptista, Robert Knott, Ian McCrone, Eleonora Po, Benjamin W. Strugnell, Katie Waine, Paul Wood, Walid T. Khaled, Huw D. Summers, Paul Rees, John W. Wills, Katherine Hughes

## Abstract

Signal Transducers and Activators of Transcription (STATs) regulate mammary gland development. Here we investigate the expression of pSTAT3 in the murine and bovine mammary gland around the day of birth. We identify polarisation of mammary alveoli towards either a low- or high-proportion of pSTAT3 positive alveolar epithelial cells. We present localised colocation analysis applicable to other mammary studies where identification, quantification and interrogation of significant, spatially congregated events is required. We demonstrate that pSTAT3-positive events are multifocally clustered in a non-random and statistically significant fashion. This finding represents a new facet of mammary STAT3 biology meriting further functional investigation.

---

The mammary gland exhibits extensive postnatal development [1-3]. Signal Transducers and Activators of Transcription (STATs) are classically activated by phosphorylation, and play key roles in regulating this development [4]. STAT3 is particularly associated with post lactational regression (involution) and there is striking up-regulation of phosphorylated STAT3 (pSTAT3) following the onset of involution [5-8]. During involution STAT3 constitutes a key regulator of cell death [5, 9, 10] and modulates the mammary microenvironment [11]. A pulse of expression of pSTAT3 protein is also observed on the day of birth in mice although this has received less focus than the prolonged activation of STAT3 accompanying involution [6].

It is well-established that there is a periparturient period of immunosuppression, and cows are very susceptible to mastitis at this time [12]. Given the susceptibility of the mammary gland to mastitis around the day of birth, and that we have previously demonstrated that during involution mammary epithelial STAT3 regulates genes associated with the acute phase response and has immunomodulatory effects [11], we considered that better understanding of the distribution of pSTAT3-expressing cells will impact understanding of the periparturient mammary immune microenvironment. We therefore sought to investigate the expression of pSTAT3 in the murine and bovine mammary gland around the day of birth.

We first examined immunohistochemical expression of pSTAT3 in murine mammary tissue from 17.5 d gestation and 2 d lactation. These time points flank the previously observed pulse of mammary pSTAT3 expression that was recorded at 0 d lactation, but not at 15 d gestation or 5 d lactation [6]. Murine mammary levels of pSTAT3 expression are extremely variable at 17.5 d gestation and 2 d lactation, with large parts of the gland exhibiting minimal pSTAT3 expression. However, where nuclear pSTAT3 expression is present, indicating activated STAT3, expression is frequently restricted to individual mammary alveoli or clusters of alveoli (Fig. 1). In view of this observation, we wished to determine whether a similar pattern of pSTAT3 expression was observed in bovine mammary tissue. The gestation length of cows is affected by breed but is approximately 279-290 d, so we examined tissue from cows between 248 d gestation and 46 d lactation (Online resource 1).

**Fig. 1.**
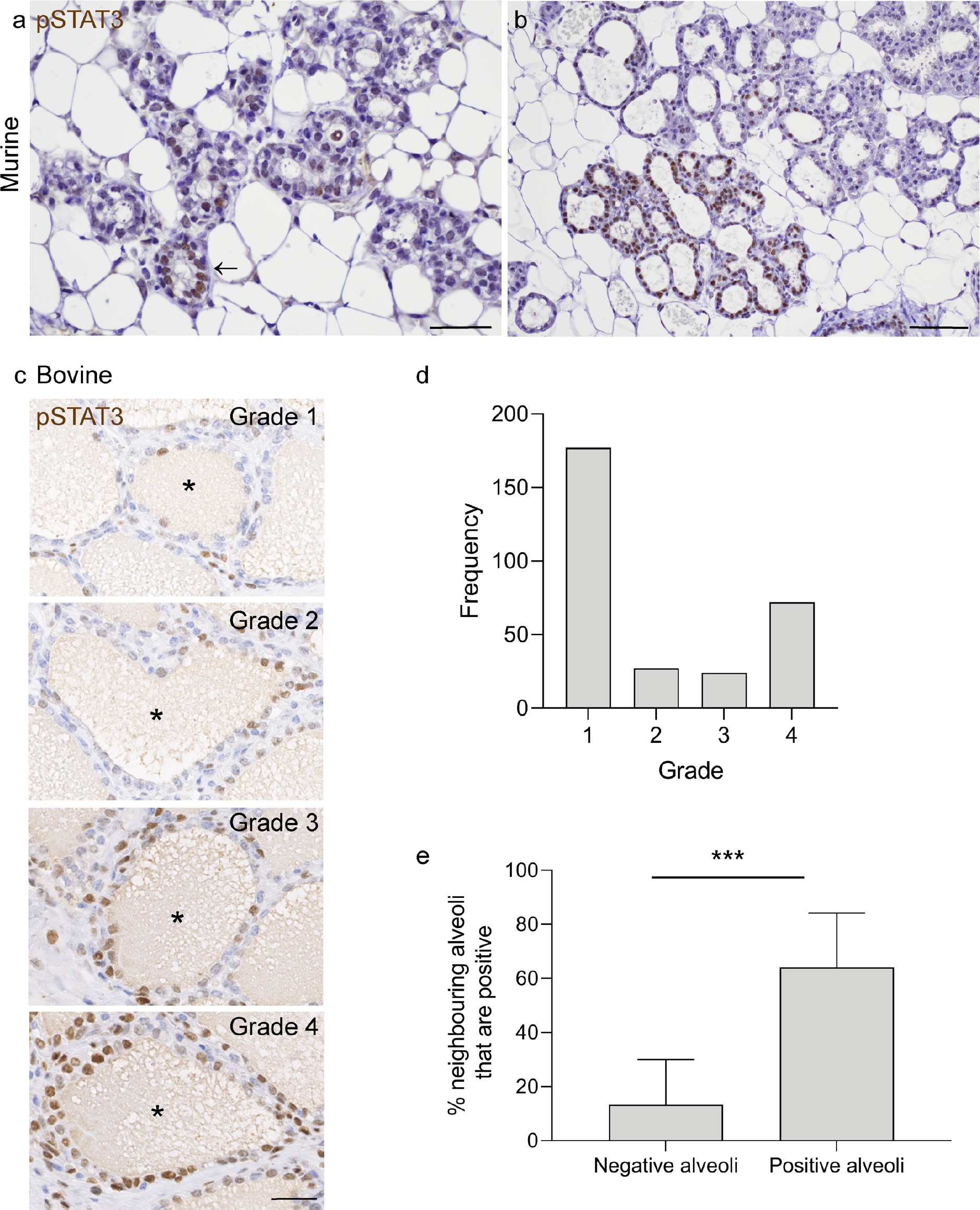
Around the day of birth there is polarisation of alveoli towards either a low- or high-proportion of pSTAT3 positive alveolar epithelial cells. (a, b) Murine tissue from 17.5 dG (a) and 2 dL (b). IHC for pSTAT3 (brown) with haematoxylin counterstain. Arrow indicates rare pSTAT3 positive alveolus. Scale bar = 40 μm (a) and 80 μm (b). (c) Example images (case 1) illustrating bovine grading scheme used to denote proportion of pSTAT3 positive alveolar luminal epithelial cells within an alveolus (*). Grade 1: 25% or less positive luminal epithelial cells; grade 2: 26 - 50% positive; grade 3: 51-75% positive; grade 4: 76 - 100% positive. IHC for pSTAT3 (brown) with haematoxylin counterstain. Scale bar = 30 μm. (d) Frequency histogram showing distribution of grades of 300 bovine alveoli selected at random from immunohistochemically stained slides from 12 mammary quarters (25 alveoli per quarter) from 7 cows. (e) pSTAT3 positive mammary alveoli have a higher proportion of pSTAT3-positive neighbouring alveoli. Results represent mean % of neighbouring alveoli that are positive for 25 alveoli analysed from each of 12 quarters from 7 cows (negative alveoli) and 7 quarters from 4 cows (positive alveoli). *** p<0.001. dG, days gestation; dL, days lactation

The udder of cows in the last third of gestation, and in early lactation, exhibits variable levels of mammary alveolar development and expansion. pSTAT3 expression levels also vary dramatically within a single mamma, between different mammae of the same animal, and between animals of similar reproductive stage. However, we noted foci with pSTAT3 expression patterns similar to those of the mouse, with pSTAT3 expression localised to individual ducts or groups of alveoli (Figure 1, Online resource 2).

Given the variable levels of alveolar expansion that we observed in the udder, and the demonstration by other investigators that a transient increase in STAT3 phosphorylation can be observed in mammary epithelial cells subjected to mechanical stress mimicking involution-associated distension [13], we considered it possible that the level of mammary alveolar dilation was affecting pSTAT3 expression. However, expression of pSTAT3 is unaffected by alveolar dimensions (Online resource 3).

Examination of tissue sections suggested that mammary alveoli are frequently composed of a majority of either pSTAT3-positive or pSTAT3-negative luminal epithelial cells. We interrogated this observation by devising a grading scheme for mammary alveolar pSTAT3 expression (Methods and Fig. 1c). We applied the grading scheme to 25 randomly selected mammary alveoli from 12 mammae from 7 individual cows, comprising a total analysis of 300 randomly selected mammary alveoli. This revealed that the frequency distribution of mammary alveolar grades is skewed towards either low (grade 1) or high (grade 4) grade and that there is a relative paucity of alveoli with a relatively even balance of cells expressing pSTAT3 and not exhibiting pSTAT3 expression (grades 2 and 3) (Fig. 1d). This indicates that in most mammary alveoli, there is a predominance of either pSTAT3-negative or pSTAT3-positive cells, and therefore suggests that there is an alveolar-level commitment to a pSTAT3 transcriptional profile.

We wished to further investigate the qualitative observation that pSTAT3-positive mammary alveoli were clustered. To analyse any potential alveolar associations, we randomly selected alveoli exhibiting any degree of pSTAT3 positivity and analysed the positivity of all adjacent alveoli in the same lobule. Mammary alveoli exhibiting any degree of luminal epithelial pSTAT3 positivity have a significantly higher proportion of pSTAT3-positive neighbouring alveoli than those mammary alveoli that are composed entirely of luminal epithelial cells in which pSTAT3 expression is not detected (Fig. 1e).

We have previously utilised spatial statistical analyses (Getis-Ord GI*) to demonstrate mammary congregation of positive immunohistochemical events [2]. In this study we performed a similar spatial analysis, adopting a local colocation quotient metric (LCQ) and further refining the image analysis by removal of the spaces created by the mammary ductular and alveolar lumina, where positive events cannot occur. This confirmed that pSTAT3-positive immunohistochemical events are multifocally clustered in a non-random and statistically significant fashion within the mammary parenchyma (Fig. 2).

**Fig. 2.**
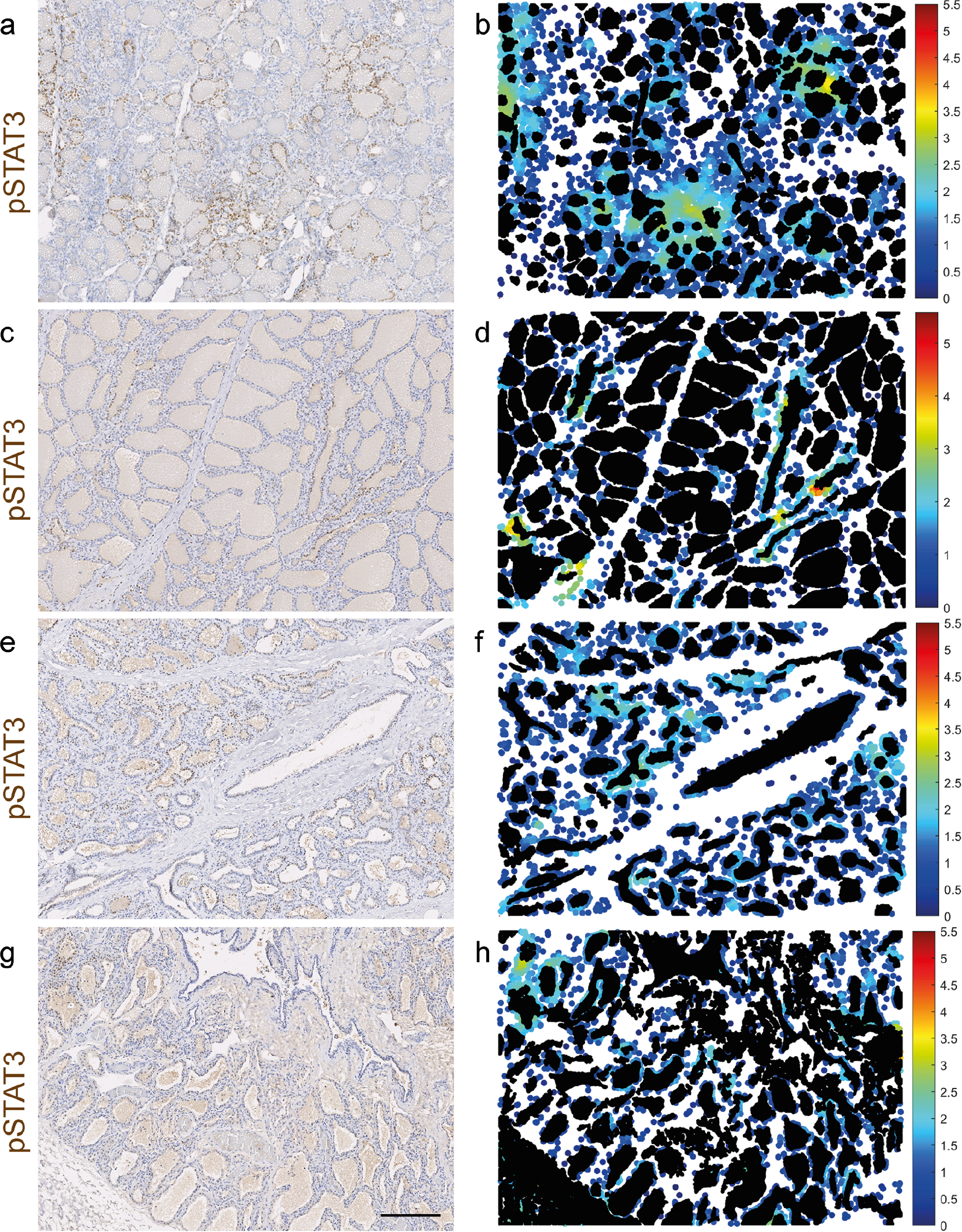
During late gestation and early lactation the bovine mammary gland exhibits hotspots of pSTAT3 expression. IHC for pSTAT3 (a, c, e, g) and accompanying spatial statistical analyses (local colocation quotient) (b, d, f, h) demonstrating regions with significant spatial congregation of pSTAT3+ cells. Mammary gland from cows 248 dG (a, b), 1 dL (c, d) 8 dL (e, f) and 46 dL (g, h); dG, days gestation; dL, days lactation. a, c, e, g Haematoxylin counterstain. Scale bar = 200 μm.

These observations shed light on an aspect of mammary STAT biology that has previously received less attention than other facets of STAT activity related to the mammary postnatal developmental cycle [4]. The finding that pSTAT3 is expressed predominantly in the mammary luminal epithelium during the periparturient period raises important questions regarding the function of this transcription factor at this developmental stage, particularly given its well-known role during post lactational regression [5, 9-11]. Although structural and functional differences between the periparturient and involution stages of mammary development may suggest a lack of commonality between these two postnatal developmental time points, a subset of STAT3 target genes that are upregulated during involution, including CD14 and leucine rich alpha-2 glycoprotein 1 (Lrg1), also exhibit upregulation on the day of birth [11, 14]. It is therefore possible that STAT3 upregulation around the day of birth may modulate the immune milieu of the gland. Given that we demonstrate multifocal hotspots of pSTAT3 expression, this may suggest that local autocrine and paracrine influences are of significance in sculpting the periparturient glandular microenvironment. This is likely important in the light of the overall susceptibility of the ruminant gland to periparturient mastitis [15].

In this analysis we capitalise on the use of spatial statistics to demonstrate significant clustering of mammary pSTAT3 expression. Assessment of the spatial distribution of cells is a powerful tool in cell biology and histopathology [16], particularly so when studying heterogeneous tissue in which marked spatial variation limits the value of whole-population statistics. Importantly, in this study the spatial analysis takes account of the structure of the mammary gland where the presence of lumina of ducts and alveoli poses a particular challenge in assessment of clustering of positive immunohistochemical events as the regions of the image not occupied by cells need to be accounted for. The localised colocation analysis presented here will be applicable to other mammary studies where identification, quantification and interrogation of significant, spatially congregated events is required.

This study has several limitations. The use of mammary tissue from non-experimental cows means that the animals sampled were of several different breeds or crosses, although the majority were Holstein Friesian cows. All the cows had concurrent morbidities, including in two cases foci of mastitis. Concurrent inflammation may influence the immune phenotype of the gland. However, in one of the two mastitis cases (case 5) both the right and left fore quarters were included in the analyses and only the left fore quarter had mastitis. In this case, both quarters had minimal pSTAT3 epithelial staining and thus the presence of an inflammatory focus appeared to have no impact. In the other case of mastitis (case 7) there was no microscopic correlation of pSTAT3 positivity with foci of inflammation. Despite the limitations of this study, the presence of clusters of pSTAT3-positive epithelial cells in the cows is strikingly similar to the tissues derived from healthy experimental mice at the defined time points of 17.5 d gestation and 2 d lactation.

This analysis raises interesting questions for future investigations, most specifically examining correlation of pSTAT3 expression with expression of STAT3 target genes. Single cell transcriptomic technologies have already been widely adopted in the mammary field [17] and spatial transcriptomics would be well suited to investigation of the role of clustered pSTAT3 expression in the periparturient gland.

Our study reveals similarities between the mouse and the cow, lending weight to the assertion that ruminants are valuable non-traditional models of mammary developmental processes [18]. Our work demonstrates that around the day of birth, in the murine and bovine mammary gland there is mammary alveolar-level commitment to a pSTAT3 transcriptional profile and pSTAT3-positive mammary alveoli are frequently grouped. pSTAT3 is an important regulator of the mammary microenvironment in other contexts. This finding therefore represents a new facet of mammary STAT3 biology meriting further functional investigation.

## Materials and methods

### Animals

Mammary tissue was collected from C57BL/6 mice at 17.5 days gestation and 2 days lactation following standard husbandry procedures. Udder tissue was collected from cows that were submitted to the diagnostic veterinary anatomic pathology service of the University of Cambridge or from cows that were euthanased by veterinarians in practice (Online resource 1). The cause of death of the animal was recorded as part of the post mortem examination procedure and/or preceding clinical investigations.

### Histology, immunohistochemistry and analyses

Mammary tissue was fixed in 10% neutral-buffered formalin. Tissues were processed using a standard methodology and 5 μm tissue sections were cut and stained with haematoxylin and eosin.

IHC followed a standard protocol using a PT link antigen retrieval system with high pH antigen retrieval solution (both Dako Pathology/Agilent Technologies, Stockport, UK). Antibody for pSTAT3 (1:100, rabbit monoclonal antibody #9145, Cell Signaling Technology) was incubated overnight at 4°C and secondary antibody was incubated for one hour at room temperature. Negative control slides were treated with isotype- and species-matched immunoglobulins.

Slides were scanned at 40× using a NanoZoomer 2.0RS, C10730, (Hamamatsu Photonics, Hamamatsu City, Japan) and were analysed with the associated viewing software (NDP.view2, Hamamatsu Photonics).

A random selection of either positive or negative alveoli were selected on a scanned slide at low magnification (25 of each per slide). Alveolar dimensions were measured using NDP.view2 and positive alveoli were assigned a grade 1-4. Grade 1 alveoli were those exhibiting 25% or less pSTAT3-positive luminal epithelial cells, grade 2 alveoli exhibited 26-50% pSTAT3-positive luminal epithelial cells, grade 3 alveoli exhibited 51-75% pSTAT3-positive luminal epithelial cells, and grade 4 alveoli were those with 76% or more pSTAT3-positive luminal epithelial cells. The number of immediately adjacent alveoli were counted and those with positive epithelial cells noted and converted to a percentage of positive neighbours for the initially selected alveolus.

### pSTAT3: Local correlation quotient spatial statistics

The centroid locations of pSTAT3+ and pSTAT3-nuclei alongside masks for tissue ‘void areas’ unpopulated by cells were extracted using pixel-classification machine learning using the freely available Ilastik and CellProfiler softwares using methods described in previous works [19] Statistically significant clustering of pSTAT3+ events relative to what would be expected by random chance were identified using the local correlation quotient (LCQ) statistic [20, 21] defined as:

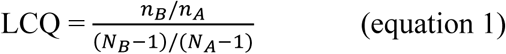

Where *N*_*A*_ is the global number of all nuclei and *N*_*B*_ is the global number of pSTAT3+ nuclei. *n*_*A*_ is the local number of all nuclei and *n*_*B*_ the local number of pSTAT3+ nuclei. The size of the local area was dynamically set for each cell according to the local cell density. A Gaussian spatial filter was used with a bandwidth equal to the distance to the 10^th^ nearest neighbour. To enable reproducibility, all image-data, image analysis steps in Ilastik and CellProfiler as well as the MATLAB code used to calculate the LCQ measure are available for download from the BioStudies database under accession number S-BSST1025 (https://www.ebi.ac.uk/biostudies/studies/S-BSST1025).

## Acknowledgements

The authors would like to thank Debbie Sabin, Ross Barker and Emma Ward of the Department of Veterinary Medicine, University of Cambridge, for their excellent technical expertise in the preparation of tissue sections, and Mathew Rhodes, of the same department, for technical assistance in the post mortem room. The authors gratefully acknowledge the Ethics and Welfare Committee of the Department of Veterinary Medicine, University of Cambridge, for their review of the study plan relating to the use of tissue from bovine post mortem examinations for the study of mammary gland biology (references: CR223 and CR625). Parts of this data were presented in oral abstract form at the Anatomical Society Summer Meeting 2023 in Bangor, Wales (25-27 July 2023; presentation date 26 July 2023).

## Statements and declarations

### Ethics approval

The Ethics and Welfare Committee of the Department of Veterinary Medicine, University of Cambridge, reviewed the study plan relating to the use of tissue from bovine post mortem examinations for the study of mammary gland biology (references: CR223 and CR625). For mouse work, all animals were treated according to local ethical committee and UK Home Office guidelines.

### Consent

Owners of animals provided informed consent for tissue to be collected for research purposes.

### Authors’ contribution statements

LJAH and KH designed the study. KH supervised the study. LJAH, BPD and KH conducted the experiments. Data was collected and/or analysed by LJAH, BPD, MB-R, CD, AFB, RK, IMC, EP, BWS, KW, PW and KH. The laboratory of WTK provided the murine tissue. Mouse experiments were conducted by SP. HDS, PR, and JWW performed the computational image analysis and spatial statistics. The first draft of the manuscript was written by KH. All authors critically revised the manuscript and read and approved the final manuscript and author contribution statements.

## Online Resources

**Online resource 1.**
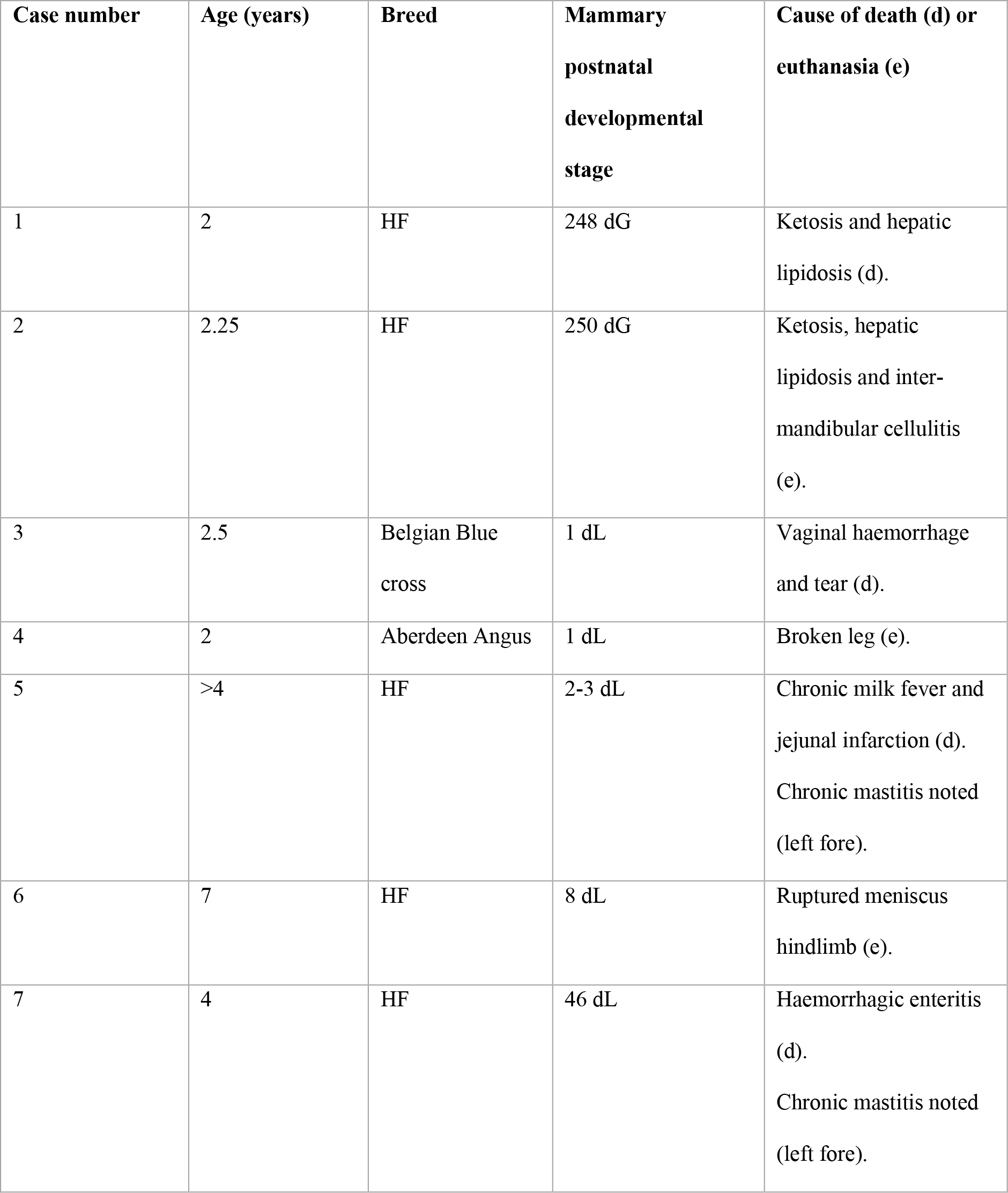
Table detailing bovine samples used in this study. dG days gestation; dL days lactation; HF Holstein Friesian.

**Online resource 2.**
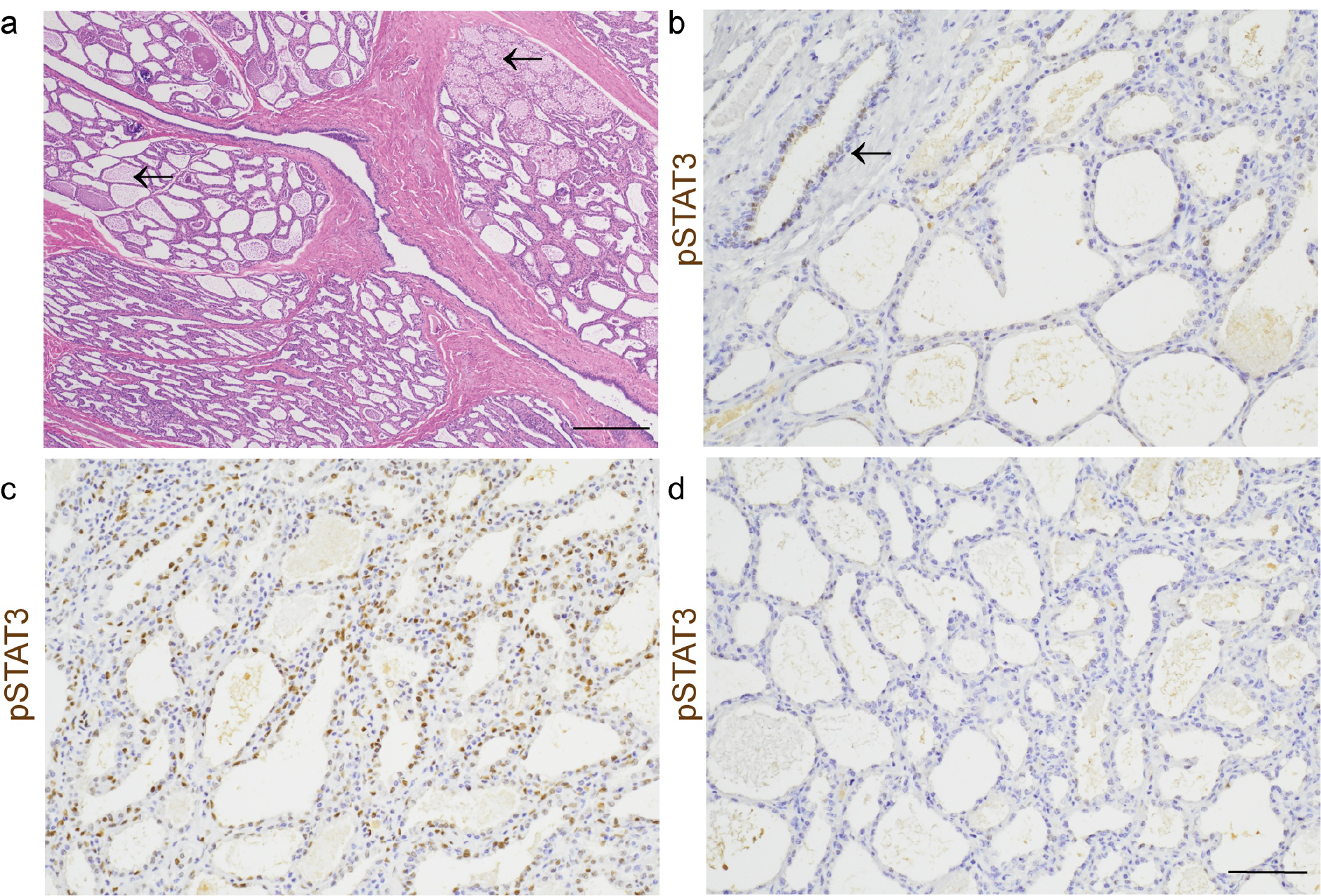
The mammary gland of cows in the last third of gestation, and in early lactation, exhibits variable levels of mammary alveolar development and pSTAT3 expression. Bovine tissue from 8 dL (case 7). (a) Alveoli are multifocally expanded with proteinaceous secretion (arrows). (b) pSTAT3 is multifocally expressed in ducts (arrow). (c) Multifocally there is intense pSTAT3 expression (c) but in other locally extensive foci there is minimal pSTAT3 expression (d). Haematoxylin and eosin staining (a) and IHC for pSTAT3 (brown) with haematoxylin counterstain (b, c, d). Scale bar = 400 μm (a) and 80 μm (b, c, d).

**Online resource 3.**
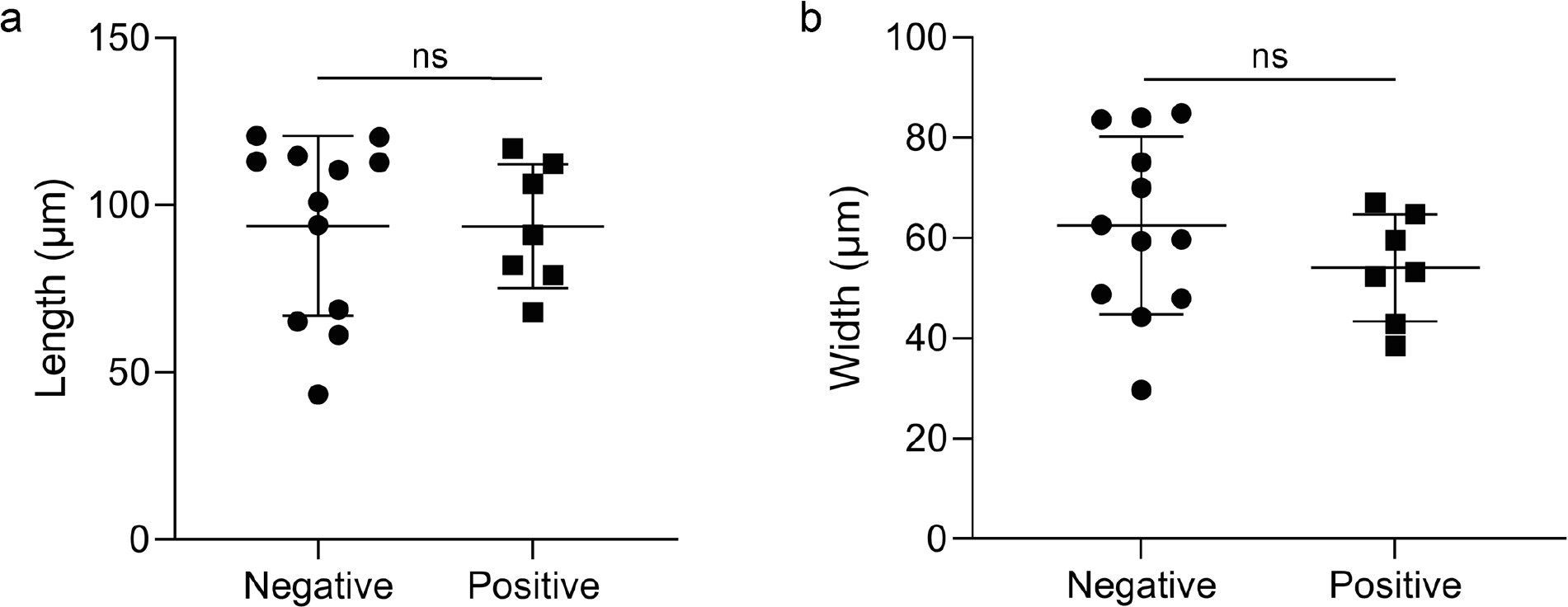
Expression of pSTAT3 is unaffected by mammary alveolar dimensions. Scatter plots demonstrating that the distribution of alveolar lengths (a) and widths (b) does not differ significantly between alveoli without any luminal epithelial pSTAT3 expression (denoted negative) and those with any degree of luminal epithelial pSTAT3 expression (denoted positive). Dots represent mean alveolar dimensions per quarter for 12 quarters from 7 cows (negative alveoli) and 7 quarters from 4 cows (positive alveoli). Bars represent mean ± standard deviation; ns, not significant.

